# SAIGE-BRUSH: an efficient, user-friendly and low cost cloud implementation for genome-wide association studies

**DOI:** 10.1101/2021.05.07.443171

**Authors:** T.M. Brunetti, N. Pozdeyev, M. Daya, K.C. Barnes, N. Rafaels, C. Gignoux

**Affiliations:** University of Colorado School of Medicine, Colorado Center for Personalized Medicine

**Author notes:** Contributed equally (N. Rafaels, C. Gignoux).

## Abstract

SAIGE-Biobank Re-Usable SAIGE Helper (SAIGE-BRUSH) allows users with little computational expertise to utilize SAIGE for GWAS with parallelization and data collection on biobank data sets. This implementation requires no installation and has additional features not programmed within the original SAIGE framework, such as concurrency, reproducibility, reusability, scalability, association analysis results filtering and output plots. This is all achieved without writing any code from the user. This implementation is currently being utilized by the Biobank at the Colorado Center for Personalized Medicine (CCPM) on Google Cloud but is flexible for a number of architectures available to genetic analysts. Availability: This open source implementation is freely available at https://github.com/tbrunetti/SAIGE-BRUSH and is licensed under the MIT License. Contact: Chris Gignoux at & Nick Rafaels at Supplemental Material: For detailed user documentation, please visit https://saige-brush.readthedocs.io/en/latest/

## Introduction

Genome-wide association studies (GWAS) are rising in popularity in part due to the emergence of new biobanks across the globe. Biobanks allow researchers to address complex genetic questions by providing large sample numbers (>20k individuals) genotyped at a relatively low cost due to improvements in data generation technology and more diverse imputation panels, which have addressed some of the criticisms and limitations of GWAS [1]. These large datasets have generated a need for better algorithms and informatics tools that can handle large, diverse data sets more efficiently while also controlling for large imbalances in binary GWAS phenotypic outcomes common in biobank-based cohorts. SAIGE is open-source software developed to address these issues while significantly reducing computational burden by generating pre-computed association parameters with a sparse kinship matrix prior to testing SNP associations with the phenotypic outcome [2]. Furthermore, SAIGE controls for case-control imbalances using saddlepoint approximation (SPA) to generate more accurate p-values [2]. Although the SAIGE framework is a commonly-used method, it relies on the end user to be able to install the software, write and submit code for analysis, and interpret and generate cleaned sets of GWAS results. To democratize and improve the utility of the software, we have developed an implementation of the SAIGE pipeline, SAIGE-<u>B</u>iobank <u>R</u>e-<u>U</u>sable <u>S</u>AIGE <u>H</u>elper (SAIGE-BRUSH), that permits a user to use SAIGE with minimal programming expertise. With one configuration file, SAIGE-BRUSH can run all aspects of analysis and provide publication-quality figures, enabling multi-trait analyses at scale.

## Implementation and Features

SAIGE-BRUSH requires no installation and is operating system agnostic, the only user software dependencies required are GoLang v>=1.13.5 [3] and Singularity v>3.0 [4]. SAIGE v.0.39 code and necessary software packages are wrapped into a Singularity container built upon a Linux operating system. Using a Singularity container allows users to move the pipeline onto any shared high performance computing system or cloud service without the security hazards of other containerization solutions [4], making the pipeline user-downloadable, fully portable and reproducible.

The novelty and strength of the pipeline is that the user never directly interacts with the container. SAIGE-BRUSH provides a pre-compiled Go binary executable that orchestrates all container interactions, pipeline logic, pipeline data management and parallelization/concurrency. A user-defined configuration file, including relevant pipeline logic, location of files, and parameters, is the only argument at runtime (see supplementary readthedocs). Within the configuration file, the user has the ability to control when and where the pipeline starts, continues, and stops without having to submit or write code at each step within the SAIGE framework; this is accomplished by simply changing pipeline logic boolean values (see Figure 1a). The configuration file allows the user to set available compute core resources, however, if left unspecified, the pipeline automatically detects the number of available cores and concurrently runs jobs as dependencies are met and resources become available. No knowledge of containerization, parallelization, or concurrency is required to use SAIGE-BRUSH efficiently.

**Figure 1.**
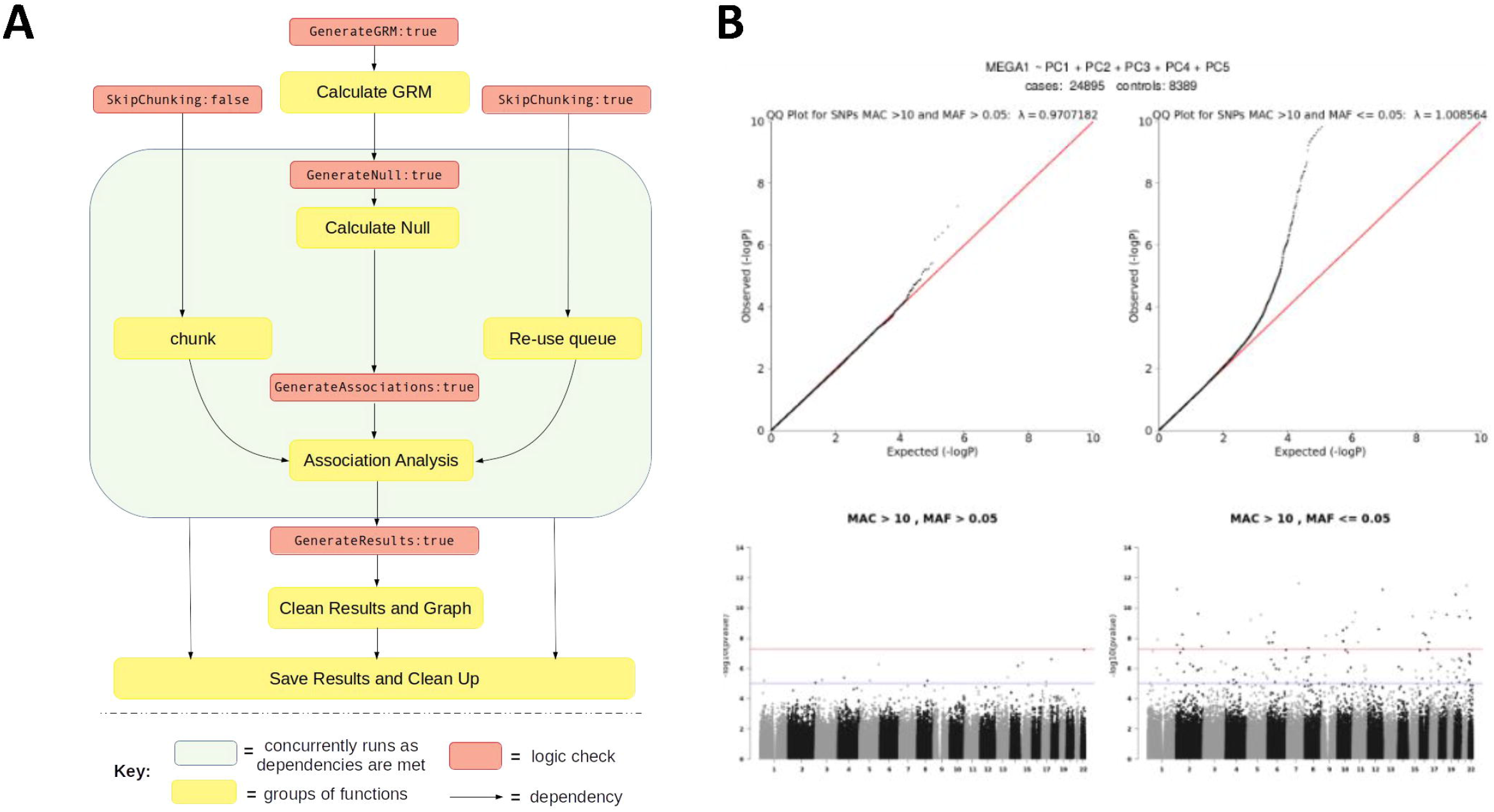
SAIGE-BRUSH implementation. (A) Schematic of implementation and logic of pipeline (B) Example of automated graphing capability

SAIGE-BRUSH includes multiple optimizations to reduce the wall time for running the pipeline. The pipeline exploits the independent nature of association analyses per variant, by generating windows along the length of user-defined input of chromosome ranges. This works for both imputed and genotyped data. Each window runs independently ensuring maximal utilization of available CPU resources. GWAS analysis of the Biobank at CCPM data containing ~ 50M of genotyped and imputed variants for 34k participants can be executed on preemptible Google Cloud virtual machines (VM) with 64 AMD-based vCPUs and 8x local SSD. When compared to chromosome or genome-wide runs taking more than 24 hours, the shorter overall runtime of SAIGE-BRUSH allows significant cost savings by use of preemptible VM pricing in Google Cloud. The entire pipeline from the Biobank at CCPM raw imputed data to QC’d results, basic summary statistics, and output graphs can be completed on average in 10.8 hours, leading to a total cost of roughly $9.30 per run. SAIGE-BRUSH allows reusing of previously computed 1M base windows when running GWAS analysis for multiple phenotypes on the same input genetic data. SAIGE-BRUSH implements additional features that are not part of the original SAIGE software. A new feature we have added is the compilation of association analysis results, clean up and visual generation of results. The pipeline automatically flips alleles to ensure that beta values are based on the minor allele, calculates minor allele counts, odds ratios, confidence intervals, and merges summary statistics with information on imputation and genotype quality. The user can define minor allele count thresholds and allele frequency thresholds to categorize variants as common or rare. These parameters are used to generate qqplots with calculated lambda inflation values [5] and Manhattan plots for common and rare variants (see Figure 1b).

SAIGE-BRUSH has been implemented as a production GWAS pipeline at the CCPM Translational Informatics Services at the University of Colorado and GWAS results have been served to dozens of investigators including population geneticists, bioinformaticians, clinicians, and basic scientists. As a user-friendly application designed to improve adoption and performance, the opportunity for this framework to improve the utility of other software packages is clear. As an open-source release, we anticipate this framework can be easily adapted in the future to other genetic software analysis frameworks, thereby reducing the learning curve and improving reproducibility of efficient GWAS at biobank scales.

## Acknowledgements

We would like to thank all of the developers of the SAIGE pipeline, of which our automated pipeline is fully based upon. Thank you to the team at Health Data Compass, Kristy Crooks, Laura Wiley, Stephen Wicks, Tzu Phang, and John Finigan for their contributions to biobank data generation, IT infrastructure and support, and domain knowledge. Additionally, we sincerely thank all participants in the Biobank at CCPM.

